# Stirred Suspension Bioreactor Culture of Porcine Induced Pluripotent Stem Cells

**DOI:** 10.1101/644195

**Authors:** Kyle Burrell, Rkia Dardari, Taylor Goldsmith, Derek Toms, Daniel A.F. Villagomez, W. Allan King, Mark Ungrin, Franklin D. West, Ina Dobrinski

## Abstract

Induced pluripotent stem cells (iPSCs) are an attractive cell source for regenerative medicine and the development of therapies, as they can proliferate indefinitely under defined conditions and differentiate into any cell type in the body. Large scale expansion of cells is limited in adherent culture, making it difficult to obtain adequate cell numbers for research. It has been previously shown that stirred suspension bioreactors (SSBs) can be used to culture mouse and human stem cells. Pigs are important pre-clinical models for stem cell research. Therefore, this study investigated the use of SSBs as an alternative culture method for the expansion of iPSCs. Using an established porcine iPSC line as well as a new cell line derived and characterized in the current study, we report that porcine iPSCs (piPSCs) can grow in SSB while maintaining characteristics of pluripotency and karyotypic stability similar to cells grown in traditional two-dimensional static culture. This culture method provides a suitable platform for scale up of cell culture to provide adequate cell numbers for future research applications involving porcine induced pluripotent stem cells.

## Introduction

Pluripotent stem cells (PSCs) are defined by two main features: the ability to proliferate indefinitely in an undifferentiated state under defined conditions, and the potential to become any cell type arising from the three germ layers, endoderm, ectoderm and mesoderm, in the body. Human and mouse embryonic stem cell (ESC) lines have significantly contributed to the progress in biomedical research and cell therapy development (1; 2), however, the ethical and safety concerns surrounding the use of human embryonic stem cells (hESCs) have limited their potential applications (3). The emergence of induced pluripotent stem cells (iPSCs) as an alternative source of cells that recapitulate the phenotype and function of ESCs, helped overcome these concerns and energized the stem cell research field (4; 5).

Outbred, large animal models that are closer in size, longevity and physiology to humans are essential to expand our knowledge gained from rodents and facilitate translation of stem cell-based approaches to human applications. iPSCs have been derived from large ungulate species including pigs that share many physiological characteristics with humans, making the pig is a powerful model for biomedical research and drug testing (6).

Large numbers of cells are required for many research and therapeutic applications, which can be challenging to obtain through conventional two-dimensional static culture systems in tissue culture plates and flasks. Two-dimensional culture systems have limited surface area, often require a large number of culture vessels and can be labor intensive. This additional handling can lead to cell loss and increased risk for contamination. Moreover, the use of numerous plates and flasks may introduce culture heterogeneity between plates and less consistency between cells. Stirred suspension bioreactors (SSBs) have been used to alleviate the problems associated with scale up of 2-dimensional static culture systems (7).

The purpose of this study was to establish an alternative, scalable culture method for the expansion of piPSCs using SSBs, based upon previous work involving human and murine PSCs (8; 9; 10; 11; 12; 13). Here we demonstrate that piPSCs from an established line and a newly generated line grown in static and SSB culture exhibit similar characteristics in terms of apparent doubling time, gene expression and differentiation potential, illustrating the utility of SSBs as an alternative culture method for the expansion of piPSCs.

## Materials and methods

### piPSC lines and culture media

The piPSCs (BT3p17) that constitutively express GFP (14) were generously donated by Drs. Telugu, Ezashi, and Roberts (University of Missouri). Briefly, these piPSCs were derived from inner cell mass cells of porcine embryos transduced with a tetracycline-inducible *hKLF4* lentiviral vector. Single factor piPSCs were further transduced using a tetracycline-inducible bicistronic lentiviral vector containing *hKLF4* and *hPOU5F1* (also known as *hOCT4)* (14). The BT3p17 line was maintained in N2B27-3i medium as previously described (15) with slight modification [1:1 ratio of DMEM/F12 (Sigma, D6421) & Neurobasal medium (Gibco, 21103049), 0.5x N2 supplement (Thermo Fisher, 17502048), 0.5x B27 supplement (Thermo Fisher, 17504044), 1000 units/mL human leukemia inhibitory factor (hLIF; Sigma, L5283), 3i cocktail [30 mM CHIR99021 (GSK3 Inhibitor; Miltenyi, 130-103-926), 40 mM PD0325901 (MEK Inhibitor; Miltenyi, 130-104-170), 10 mM PD173074 (FGFR3 Inhibitor; Sigma, P2499)], 0.5x Glutamax (Gibco, 35050-061) supplemented with 0.1 mM *β*-mercaptoethanol (Sigma, M3148), 1x MEM non-essential amino acids (MEM NEAA; Gibco, 11140050), 0.01% bovine serum albumin (Sigma, A7906), and 2 μg/mL doxycycline (DOX) (Sigma, D9891). We will refer to these cells as LIF-dependent piPSC line.

To test the utility of SSB culture on another cell line, a second piPSC line was generated for this study. Briefly, the cells were derived from dermal fibroblasts of a 2-year-old Hampshire pig (Kewanee Farm, Dudley, GA, USA). The cells were transduced using viPS lentiviral vectors (Thermo Scientific, West Palm Beach, FL, USA) that express human *OCT4, NANOG, SOX2, LIN28, KLF4* and *C-MYC* open reading frames under the control of the human elongation factor-1α (EF-1α) promoter in the presence of 1.2% GeneJammer (a polyamine-based transfection reagent; Stratagene, La Jolla, CA, USA). Cells were maintained in mTeSR™1 (STEMCELL Technologies, 85850), a serum-free medium designed for the feeder-free culture of hESC and hiPSC. The mTeSR™1 contains, among other factors, recombinant human basic FGF and recombinant human TGFβ. We will refer to these cells as LIF-independent piPSC line.

### piPSC lines culture

Cryopreserved LIF-dependent piPSCs were thawed in a 37°C water bath and washed in prewarmed medium. piPSCs (10^5^ cells/mL) were plated onto 6-well culture plates (Corning, 353846) coated with 50 μg/mL poly-d-lysine (PDL; Sigma, P0899), 20 μg/mL laminin (LN; Sigma, L-2020) and irradiated mouse embryonic fibroblasts (MEFs; ES Cell Facility, Center of Genome Engineering, University of Calgary) and cultured in N2B27-3i medium. piPSCs were cultured for 2 passages (6 days), with a full medium change at day 2 of 3 day passage. piPSCs were then passaged feeder-free (PDL/LN) in N2B27-3i medium, with a full medium change at day 2 and passage at day 3 using Accutase (STEMCELL Technologies, 07920).

LIF-independent iPSCs were cultured on mitomycin-treated feeder cells for several passages, and then individual colonies were transferred onto Matrigel-coated plates (Corning, 354277) and characterized. Cells originating from a single colony were included in this study. The LIF-independent piPSC line was maintained in mTeSR™1 medium at 37°C, 5% CO_2_, in saturated humidity, and passaged using Accutase, when cell confluency reached 80%. In all culture conditions, piPSCs were dissociated into single cells, counted and assessed for cell viability using a hemocytometer and Trypan Blue exclusion (Sigma, T8154).

Both piPSC lines were cultured to obtain adequate starting cell numbers for comparative culture experiments, gene expression analysis, differentiation assays and cytogenetic analysis (these cell populations are referred to as pre-culture piPSCs or pre-piPSCs. The experimental design is schematically outlined in Fig. 1.

**Fig 1.**
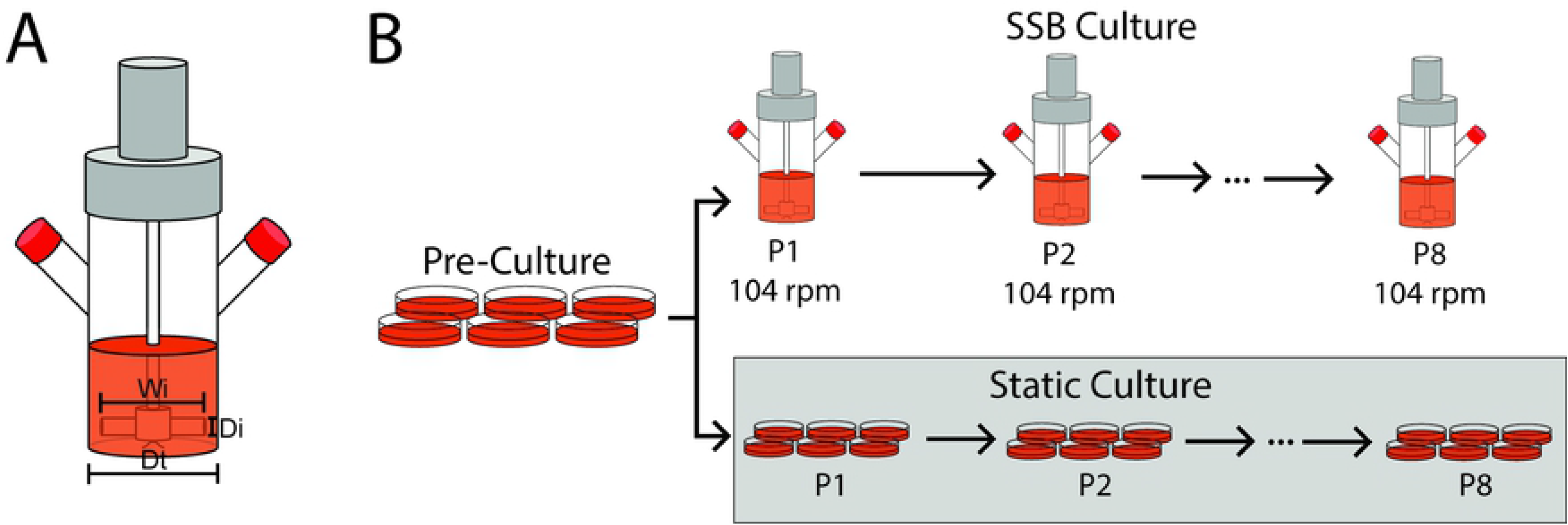
Experimental design. Schematic representation of SSB, where Wi, Di, Dt represents an impeller width of 0.95 cm, an impeller diameter of 3.5 cm, and a vessel diameter of 3.88 cm, respectively (A). Schematic representation of comparative culture (B).

### piPSC static culture

For static culture, 10^5^ cells/mL from the initial pre-piPSC culture of LIF-dependent and LIF-independent cells were plated onto 6-well culture plates coated with PDL/LN and Matrigel-coated plates, respectively, with a full medium change at day 2 and passage at day 3 using Accutase. Each subsequent passage used piPSCs generated from previous static culture up to passage 8.

### piPSC SSB culture

For SSB culture, 50 mL magnetically driven suspension bioreactors (NDS Technologies, Inc.; Spinner Flask Assembly Complete, 264500-50) were siliconized as per manufacturer’s instructions (Sigma, SL-2) and used with a working volume of 50 mL (Table 1, Fig 1). Fifty mL SSBs were inoculated with 1.406 x10^6^ piPSCs from the pre-piPSC culture resuspended in 50 mL N2B27-3i medium or mTeSR™1. The piPSCs were agitated at 104 rpm, corresponding to a maximum shear stress of 3.0 dyne/cm^2^. The cells were passaged on day 3 using Accutase. Subsequent SSB cultures were inoculated using piPSCs generated from the previous SSB culture up to passage 8.

**Table 1.** 50mL SSB and medium parameters.

| Parameter | Value | Units | Symbol |
| --- | --- | --- | --- |
| Working Volume | 50 | mL | VL |
| Impeller Width | 0.95 | cm | Wi |
| Impeller Diameter | 3.5 | cm | Di |
| Vessel Diameter | 3.88 | cm | Dt |
| *\*Medium Density* | *1.006* | *g/mL* | *$\rho$* |
| *\*Medium Viscosity* | *0.85* | *cP* | *$\mu$* |
| *\*Medium Kinematic Viscosity* | *0.008449* | *cm<sup>2</sup>/s* | *$\nu$* |

\* denote values used from previous study (16).

### Apparent doubling time calculations

The piPSCs collected from the static culture and the SSB culture were counted at 72 hrs after the start of each passage. Apparent doubling times were calculated using Trypan Blue exclusion and the formula (14):

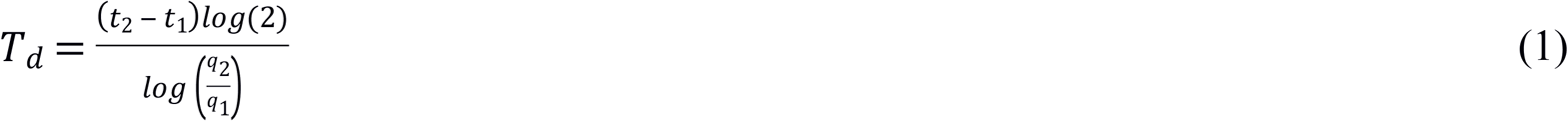

where T_d_ represents apparent doubling time, q_1_ is the initial viable cell number at t_1_, and q_2_ is the final viable cell number at t_2_ (72 hrs). t_1_ (0 hr) represents the initial inoculation time. Live cell counts were used for this calculation.

### Cell viability calculation

Live and dead cells were counted, and cell viability was determined using the following formula:

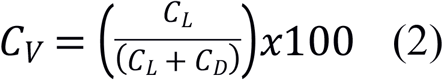

where C_V_ represents cell viability as a percentage, C_L_ is the total cells counted after 72 hrs that did not stain with Trypan Blue and are live, C_D_ is the total cells counted 72 hrs after the start of every passage that did stain with Trypan Blue and are dead.

### Shear stress calculation

Parameters, variables and equations to calculate the maximum shear stress that the piPSCs may experience during SSB culture are listed in table 1 and the following equations were used: The maximum shear stress (*τ*_max_) that a single cell is exposed to, on the surface of an aggregate can be estimated using the following equations (17):

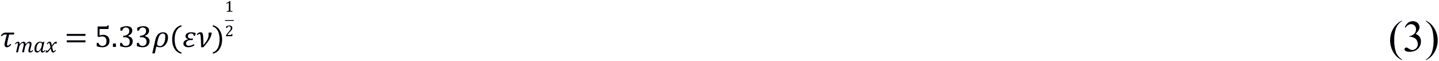

Where power dissipated per unit mass (*ε*) is calculated using the following equation:

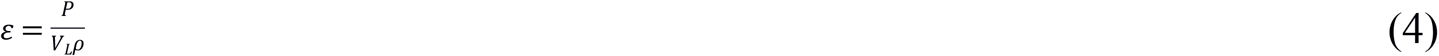

*P* is representative of the consumed power (W) and can be estimated using the equation below (18):

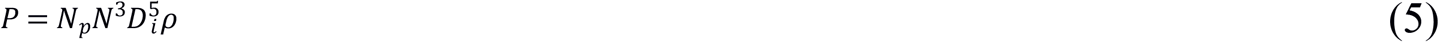

Whereas the power number (*N_p_*) can be estimated through the following equations (18):

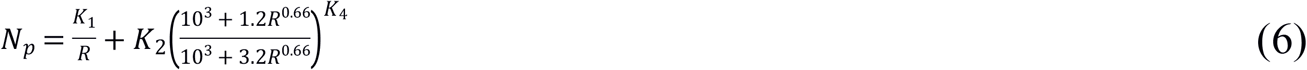

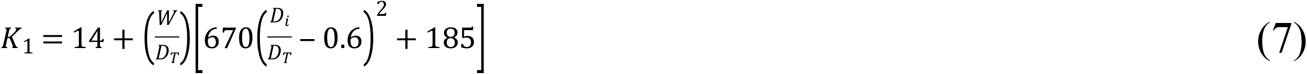

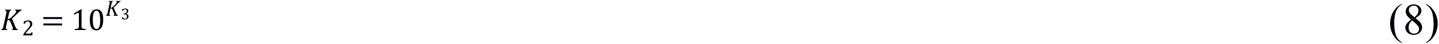

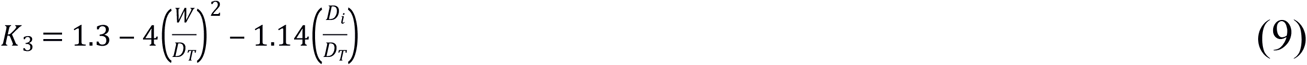

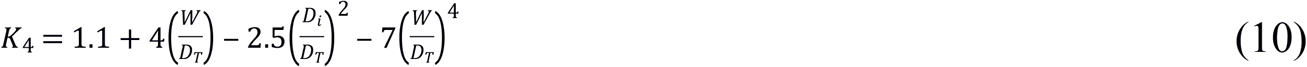

The Reynolds number (Re) can be calculated using the following equation (16):

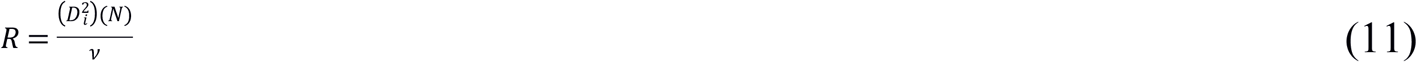

### Immunocytochemistry analysis

piPSCs were collected from pre-culture, passage 8 (P8) static and P8 SSB cultures, washed and fixed in 2% paraformaldehyde in DPBS for 20 min at room temperature, and immobilized onto cytospin slides by centrifugation at 1000 × g for 5 min., followed by permeabilization with 0.1% Trition-X (Millipore Sigma, 9410) in PBS for 10 min. Permeabilized cells were washed in PBS, blocked with CAS block (Life Technologies, 008120) for 15 min at RT and incubated with primary antibody *(SI Materials and Methods)* at 4°C overnight. After overnight incubation, cells were washed and incubated with secondary antibody *(SI Materials and Methods)* for 1hr at RT in darkness. Slides were washed and VECTASHIELD mounting medium with DAPI (Vector Laboratories, H1200) was added to slides before visualization.

### Flow cytometry

piPSCs from pre-culture, P8 static and P8 SSB cultures fixed in 2% paraformaldehyde in DPBS for 20 min at room temperature were prepared by permeabilization with 0.1% Triton-X in PBS for 10 min. Permeabilization was not required to detect the surface antigen SSEA-4. Cells were washed and blocked for 15 min with CAS block, followed by primary antibody or isotype incubation for 25 min at RT. Cells were washed and incubated with secondary antibodies for 20 min at RT in darkness. Cells were washed, resuspended in 400 μL of 1% BSA in PBS and transferred to 5 mL round-bottom FACS tubes for analysis (FACSCalibur, Becton Dickinson).

### Spontaneous differentiation in embryoid bodies

LIF-dependent piPSCs from pre-culture, P8 static and P8 SSB cultures were collected as single cells, transferred into differentiation medium [piPSC medium supplemented with 20% FBS and without LIF, 3i cocktail; with and without DOX] and plated onto 0.2% gelatin-coated (Sigma, G1393) 12-well plates (Greiner Bio-One, 665180) at an inoculation density of 1 x10^5^ cells per well. Spontaneously formed embryoid bodies (EBs) were collected after 3 days.

LIF-independent piPSCs from pre-culture, P8 static and P8 SSB cultures were collected as single cells in low attachment plates and kept in mTeSR™1 medium for 24h to allow aggregates formation. Then DMEM/F12 (Sigma, D6421) supplemented with 20% Knockout serum (KSR) medium (ThermoFischer, 10828010) was added to the formed aggregates to support differentiation, and changed every 48h. EBs were collected after 10 days of culture.

The samples from both types of cells were subjected to immunocytochemistry *(SI Materials and Methods)* and RNA extraction for gene expression analysis.

### RNA extraction and cDNA synthesis

piPSCs from pre-, static, SSB culture, and spontaneously formed EBs were collected in DPBS and transferred into 1.5 mL centrifuge tubes. The samples were then centrifuged at 1000 × g for 5 min to pellet cells. After the DPBS was removed, the cell pellets were snap-frozen in liquid nitrogen and stored at −80°C until RNA extraction. RNA was collected from piPSCs (Qiagen RNeasy Mini Kit, 74106) and differentiated cells (Qiagen RNeasy Micro Kit, 74004) as per manufacturer’s protocol. The RNA concentration was measured using a Nanodrop 2000 spectrophotometer (Thermo Scientific) and this concentration was used for cDNA synthesis. cDNA synthesis was performed via manufacturer’s instructions (Applied Biosystems High Capacity cDNA Reverse Transcription Kit; Thermo Fisher, 4368814). cDNA was made at 0.1 ng/μL for primer testing and standard curve generation and 0.01 ng/μL for qPCR.

### Quantitative Polymerase Chain Reaction (qPCR)

qPCR was performed on cDNA generated from piPSCs, spontaneous EBs, and pig fetal fibroblasts (PFF) to assess relative gene expression of pluripotency associated genes *SOX2, OCT4, cMYC, KLF4, NANOG, TERT* as well as genes specific for the three germ layers *[NESTIN* (ectoderm), *ACTA2* (mesoderm), and *GATA4* (endoderm), *GATA6* (endoderm) and *SOX17* (endoderm)] to *GAPDH or HPRT* using Applied Biosystems 7500 Fast Real-Time PCR System (Holding stage: 95 °C 20 sec; Cycling stage: 95 °C 3 sec, 60 °C 30 sec (40 cycles); Melt Curve stage: 95 °C 15 sec, 60 °C 1 min, 95 °C 15 sec, 60 °C 15 sec). Fold expression was calculated using ΔΔCt followed by a log2 transformation. Primer sequences are listed in supplementary information (Table S2). For pluripotency-associated genes, analysis was performed on piPSCs from each culture at passage 8, using pre-culture piPSCs as a control. For genes associated with differentiation, pre-culture piPSCs were used as the control to determine upregulation of differentiation-associated genes across all conditions.

### Teratoma assay

Two million piPSCs from static and SSB culture were collected as single cells, mixed with matrigel (100%) and injected subcutaneously into 5-6 week old NOD-SCID (non-obese diabetic severe combined immunodeficient) mice (Charles River). Masses were collected at 8 weeks postinjection and fixed in 4% paraformaldehyde (Ricca Chemical Company, 1120-32). Samples were processed for histological analysis. Animal procedures were approved by the University of Calgary Animal Care Committee in compliance with the Guidelines of the Canadian Council of Animal Care.

### Cytogenetic analysis

piPSCs from pre-culture, P8 static and P8 SSB cultures were cultured onto T-25 flasks (Grenier Bio-One, 690-170) coated with PDL/LN or Matigel at an inoculation density 6 x10^5^ cells in 4 mL of N2B27-3i medium or mTeSR™1 for 2-3 days until piPSCs reached 60-70% confluency. T-25 flask cultures were then cell-cycle synchronized by adding Methotrexate (10^−7^ M final concentration), for about 16 hrs of incubation in fresh cell culture media at 37.5°C. Cell replication block was released by washing the cells twice with fresh media and then incubating the cells in media containing a solution of Bromodeoxyuridine (BrdU: 1.25 mM final concentration) for 5 hrs. For chromosome preparations, cells were treated for 30 min with Colcemid (10 mg/mL), inducing metaphase arrest and consequently harvested, treated with a hypotonic solution (75 mM potassium chloride) and fixed with Carnoy’s solution (3:1 Methanol: Acetic Acid). Chromosome preparations were subjected to conventional GTG-banding protocol and standard karyotypes were obtained and analyzed as previous described (19)..

### Statistical analysis

Multiple t-tests with *post-hoc* Bonferroni multiple comparison correction and ANOVA with *post-hoc* Tukey’s multiple comparison test were used to determine statistical significance between culture groups as well as between treatments within each culture type. Significance was determined at p<0.05. Gene expression was calculated as fold change, expressed as ΔΔCt followed by a log2 transformation. Each replicate was collected from an independent cell population; aliquots of piPSCs were thawed and used for each replicate. Analyses were performed using Prism7 (GraphPad).

## Results

In the absence of any pre-established SSB culture conditions for piPSC, a protocol developed for mESC was used as a starting point [8]. Preliminary experiments testing different rotation speeds and addition of microcarriers indicated that piPSCs formed aggregates rather than attached to microcarriers. A shear stress of 3.0 dyn/cm^2^ was chosen as it supported high cell viability in piPSC aggregates. This shear stress maintained pluripotency in mESCs comparable to 6.0 dyn/cm^2^ in the presence of LIF, without cell loss due to necrotic centers in cell aggregates (9). Therefore, we used the LIF-dependent piPSC line as a first target to evaluate these culture conditions.

### Static and SSB cultured-iPSCs maintain similar cell growth rates and stable karyotype

Since piPSCs have never been exposed to the dynamic conditions observed with SSB culture, a 28 day comparative culture was designed to compare the growth of LIF-dependent piPSCs in SSBs to the static culture. Our first objective was to determine if the dynamic SSB environment changed apparent doubling time, cell viability and karyotype stability of piPSCs when compared to the static culture system (Fig. 2). Aside from passage 1 (Fig. 2A; p<0.05), there was no difference between culture conditions at any passage when considering apparent doubling times (Fig. 2A) nor overall cell viability (Fig. 2B).

**Fig. 2.**
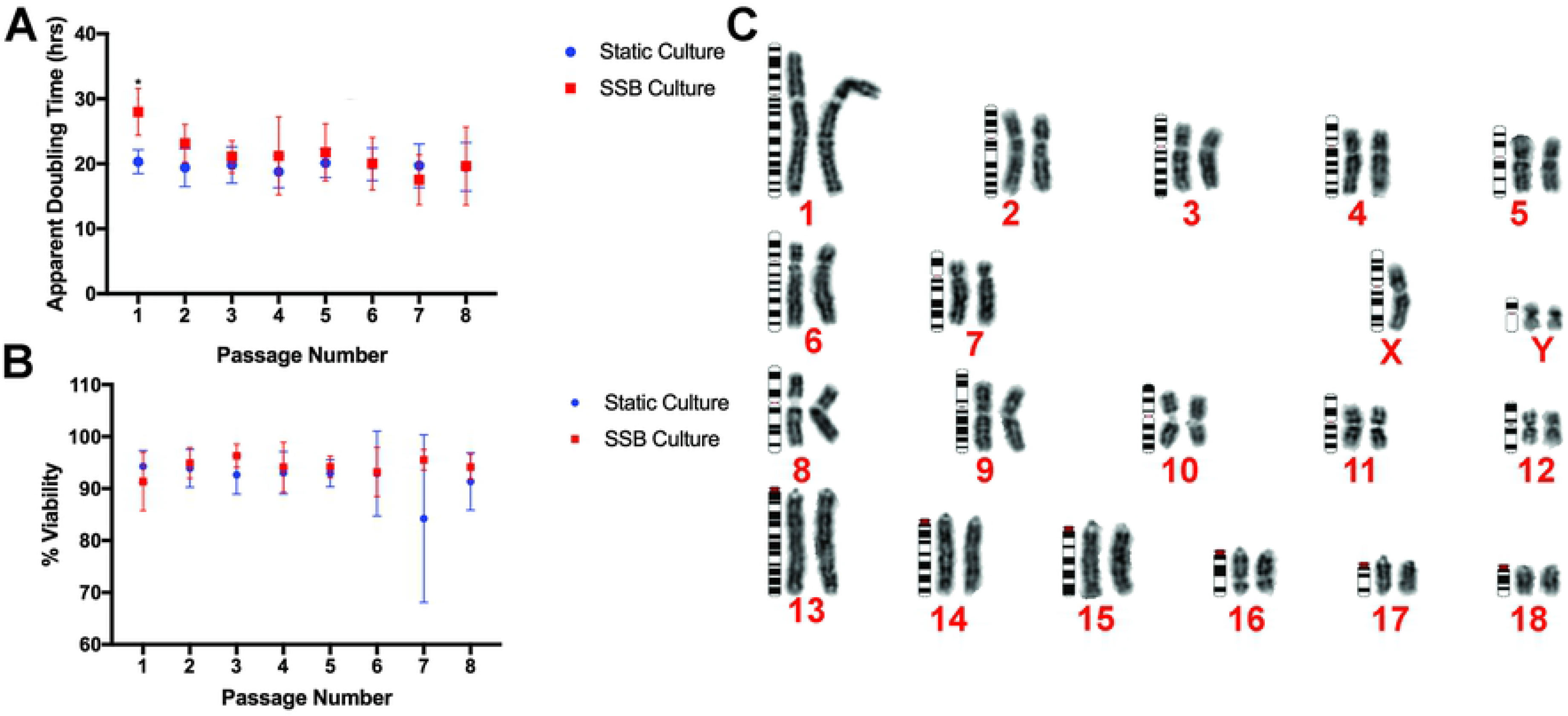
Apparent doubling time, viability and cytogenetic analysis. Apparent doubling (A) and cell viability (B) comparing SSB to static cultures over 8 passages. Cells were passaged every 3 days, and cells were collected at day 3 for analysis n=5. Error bars= SD. Cytogenetic analysis of piPSCs (C; image shows piPSCs from SSB culture). Cytogenetic analysis was performed on preculture piPSCs, P8 SSB piPSCs, and P8 static piPSCs; n=5.

Cytogenetic analysis showed a stable karyotype (>95%) throughout the duration of the culture for pre-culture, static culture, SSB culture (Fig. 2C; visual representation using piPSCs cultured in SSB culture). Each sample shows no evidence of any structural chromosome abnormality, however, all cell cultures had a 2n= 39, XYY karyotype constitution (disomy of the Y sex chromosome: Fig. 2C). Freshly thawed cells showed a similar karyotype, including disomy of the Y sex chromosome.

Samples were collected from SSB culture at each passage to visualize aggregate formation in suspension culture. Aggregates formed freely after inoculation of piPSCs as single cells (Fig. 3 B^I^ & C^I^, respectively) while maintaining GFP expression (Fig. 3 B^II^ & C^II^, respectively). piPSCs cultured statically also maintained GFP expression (Fig. 3 A-A^II^).

**Fig. 3.**
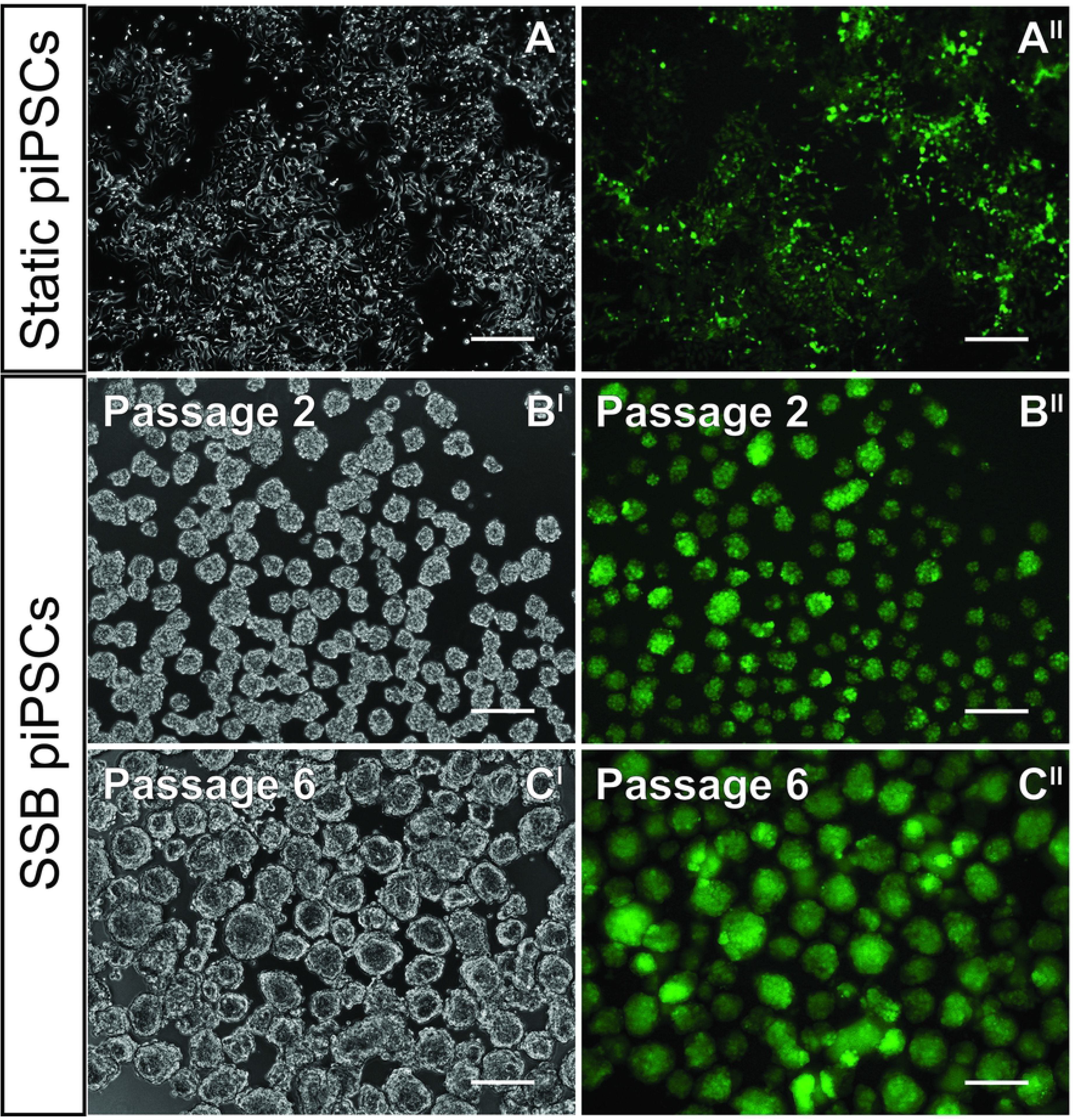
Appearance of piPSCs in static and SSB culture. piPSCs (constitutively expressing GFP) grown in static (A & A^II^) and SSB (B^I^, B^II^, C^I^, C^II^). Aggregates formed in SSB culture are of relatively uniform size (61.8 ± 1.6 μm in diameter at P6 and 87.5 ± 14.2 μm at P8, n=462 aggregates) and shape; scale bar 200 μm.

### SSB cultured-iPSC line maintains the expression of pluripotency markers

In order to determine if dynamic culture conditions affect the maintenance of pluripotency in piPSCs, we assessed the expression of pluripotency-associated genes by qPCR, flow cytometry and immunocytochemistry. Pre-culture piPSCs were used as a control to identify any differences observed in gene or protein expression attributed to long term culture. No statistically significant difference was detected by qPCR in expression of *SOX2, OCT4, KLF4*, and *cMYC* between cells from pre-, static, and SSB culture conditions (Fig. 4A). The similarity of piPSCs from static and SSB cultures was also apparent using flow cytometry analysis, which determined that there were no significant differences in the percentage of cells expressing the pluripotency-associated markers SSEA-4, SOX2, and OCT4, as well as the proliferation marker Ki67 (Fig. 4B). These similarities were qualitatively confirmed by immunocytochemistry for SSB-cultured piPSCs (Fig. 4C) and piPSCs cultured statically (Fig. 4D).

**Fig 4.**
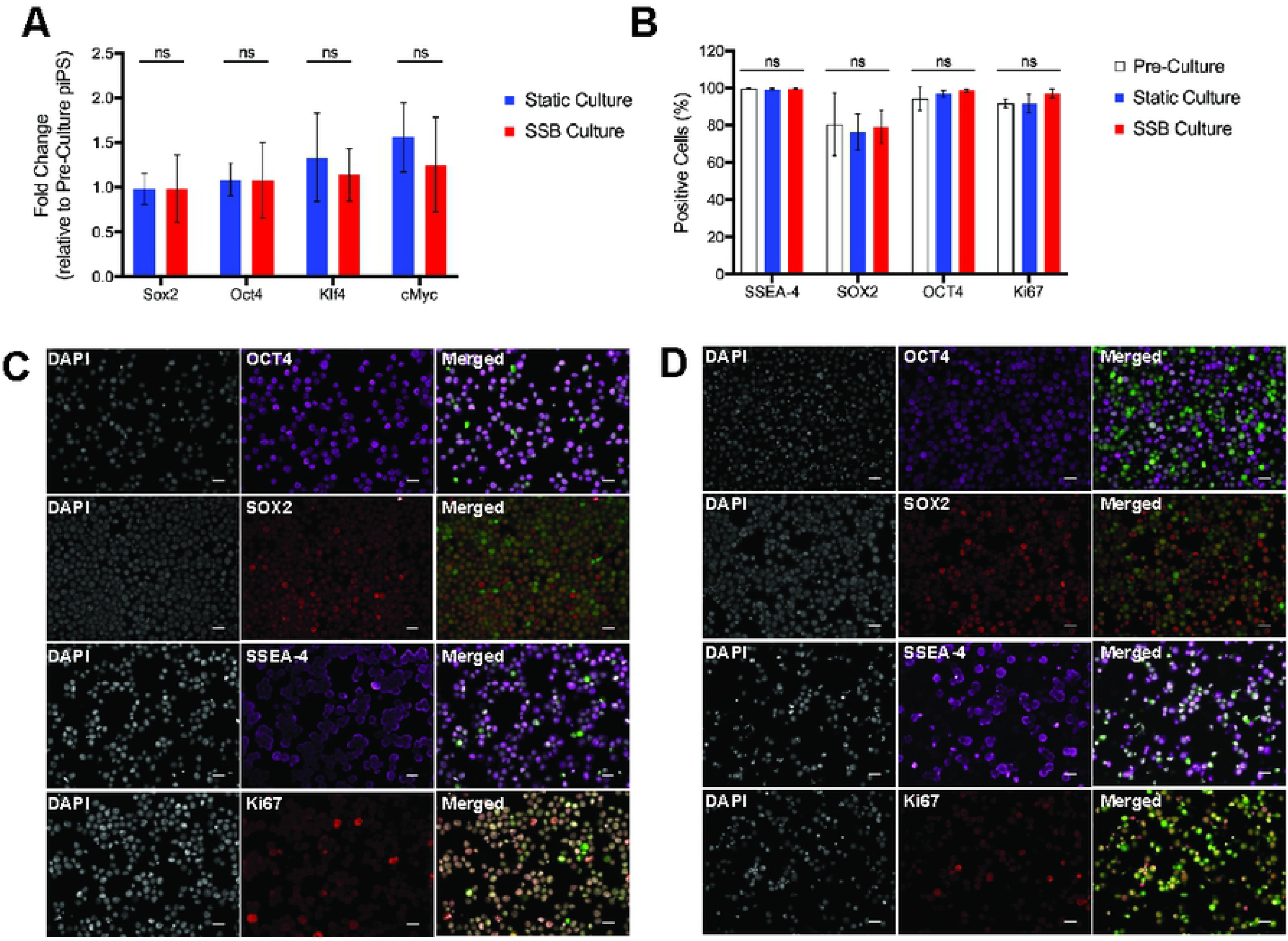
Protein and gene expression of pluripotency-associated and proliferation markers. **A**: qPCR analysis assessing relative gene expression levels of pluripotency-associated markers; n=4, **B:** Flow cytometry illustrating percentage of pluripotency-associated (OCT4, SOX2, SSEA-4) and proliferative (Ki67)-positive piPSCs; n=3, **C,D:** Immunocytochemistry of piPSCs cultured in SSBs **(C)** and piPSCs cultured statically **(D)** for pluripotency-associated (OCT4; purple, SOX2; red, SSEA-4; purple) and proliferation (Ki67; red) markers; scale bar 50 μm.

As controls, isotype controls and unstained piPSCs were used to validate staining of piPSCs from pre-, static, and SSB culture conditions (Fig. S1).

### SSB cultured-iPSC line differentiates into three germ layers in *vitro*

The ability to differentiate into derivatives of all three germ layers is a characteristic of pluripotency. In order to assess the differentiation potential of the piPSCs *in vitro*, differentiation assays were performed. The LIF-dependent piPSCs used in this study were reprogrammed to a pluripotent state using a tetracycline-inducible lentiviral vectors [14], and the continued expression of these lentiviral vectors depends on presence of tetracycline, or DOX, in the culture medium. Beside its role in controlling the expression of pluripotency markers in DOX-inducible iPSCs, DOX has been proven to trigger phosphoinositide 3-kinase-Akt signaling pathways involved in maintaining survival and promoting the self-renewal of ESC and iPSC (20). Upon removal of DOX, cells are expected to differentiate. The assessment of the differentiation potential of iPSC in the presence of DOX will address the question whether DOX alone is able to prevent the differentiation iPSC by maintaining the expression of pluripotency markers or additional factors present in 3i serum free medium are required to support such an effect.

After 3 days of spontaneous differentiation in differentiation medium, embryoid bodies (EBs) were assessed for gene and protein expression of NESTIN (ectoderm), ACTA2/*α*SMA (mesoderm), and GATA6 (endoderm). A marked upregulation of *ACTA2* (Fig. 5A; a-b: p<0.05; a-c: p<0.05; b-c: p<0.05; n=5), *GATA6* (Fig. 5B; a-b: p<0.05; n=5), and *NESTIN* (Fig. 5C; a-b: p<0.05; a-c: p<0.05; b-c: p<0.05; n=5) expression was observed in differentiated pre-, static and SSB piPSCs. As expected, upregulation of differentiation associated genes was more pronounced when comparing undifferentiated piPSCs with piPSCs differentiated without DOX. Although *NESTIN* expression differed between control pre-culture piPSCs and the piPSCs differentiated in the presence of DOX, qPCR analysis showed that the expression levels of *GATA6* and *ACTA2* were not significantly different between these cell populations. The piPSCs used in this study were reprogrammed to pluripotent state using tetracycline-inducible lentiviral vectors (15), and the continued expression of these lentiviral vectors depends on presence of tetracycline, or DOX, in the culture medium. This explains more robust differentiation of the piPSCs upon the removal of DOX in addition to removing the 3i cocktail, LIF, and the addition of 20% FBS.

**Fig 5.**
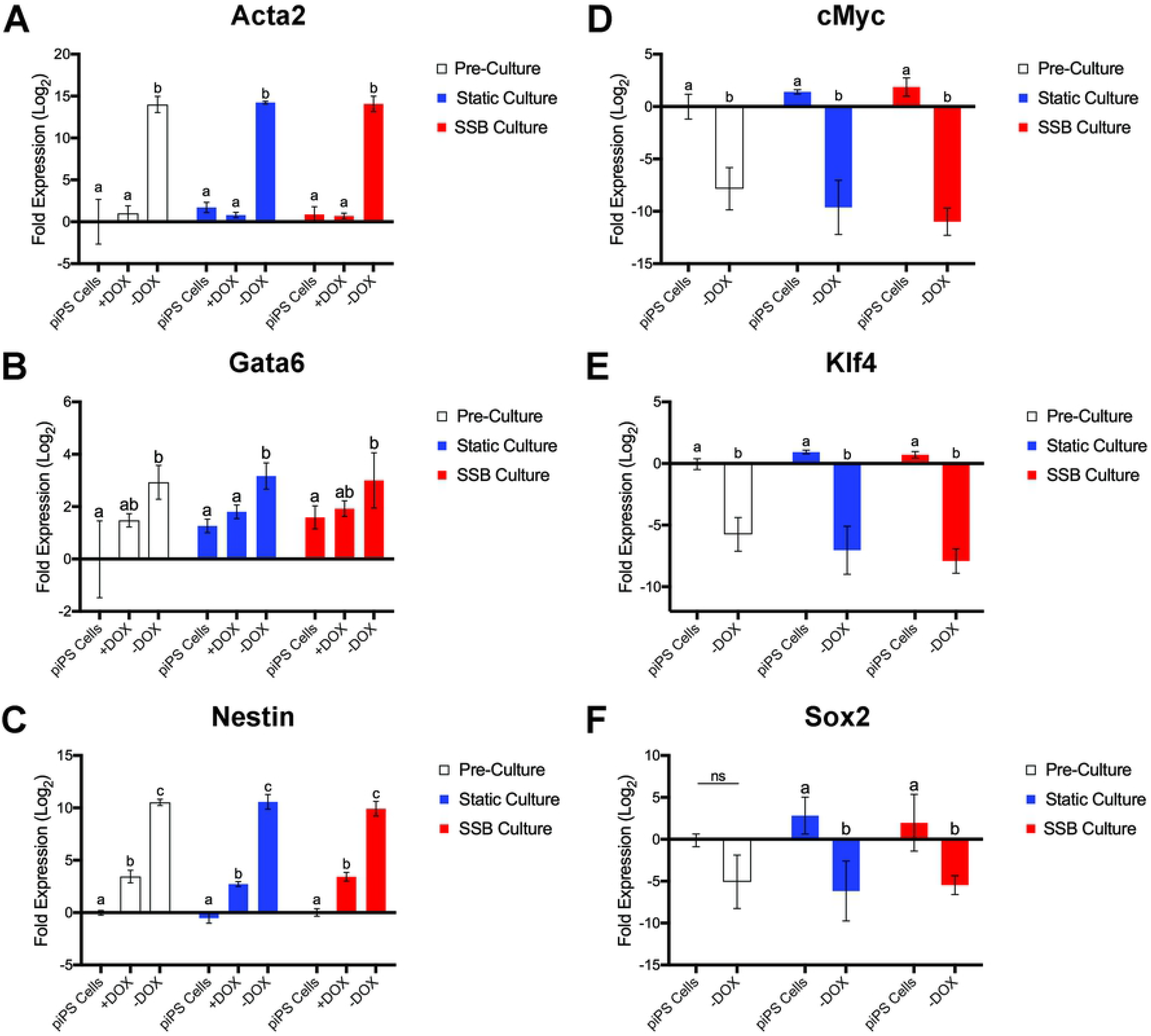
Gene expression analysis. Upregulation of genes indicative of differentiation (A-C) and the subsequent downregulation of pluripotency-associated gene expression (D-F) on day 3 of spontaneous embryoid body differentiation. Differentiation medium with or without DOX was used to determine the differentiation capacity of the piPSCs. Significant differences in gene expression were observed for A) *ACTA2* (a-b: p<0.05; a-c: p<0.05; b-c: p<0.05); B) *GATA6* (a-b: p<0.05); C) *NESTIN*(a-b: p<0.05; a-c: p<0.05; b-c: p<0.05); D) *cMYC* (a-b: p<0.05); E) *KLF4* (a-b: p<0.05); F) *SOX2* (a-b: p<0.05). Gene expression was normalized to *GAPDH* and represented in log2 scale; n=5. Error bars= SD.

At the same time, analysis of the expression of pluripotency-associated genes demonstrated downregulation of *cMYC* (Fig. 5D; a-b: p<0.05; n=5), *KLF4* (Fig. 5E; a-b: p<0.05; n=5), and *SOX2* (Fig. 5F; a-b: p<0.05; n=5) upon differentiation in the absence of DOX, (Fig. 5C). Expression of *OCT4* was not analyzed under these conditions. Downregulation of pluripotency-associated genes is consistent with the data demonstrating upregulation of differentiation-associated genes as pluripotency-associated genes are expected to become downregulated upon differentiation to lineage specific cells. Quantification of the differentiation- and pluripotency-associated gene expression is relative to pre-culture piPSCs, which were used as a control.

Supporting the gene expression data, lineage specific protein expression was observed by staining for *α*SMA and NESTIN in piPSCs from pre-, static, and SSB cultures allowed to spontaneously differentiate for 3 days (Fig. 6).

**Fig 6.**
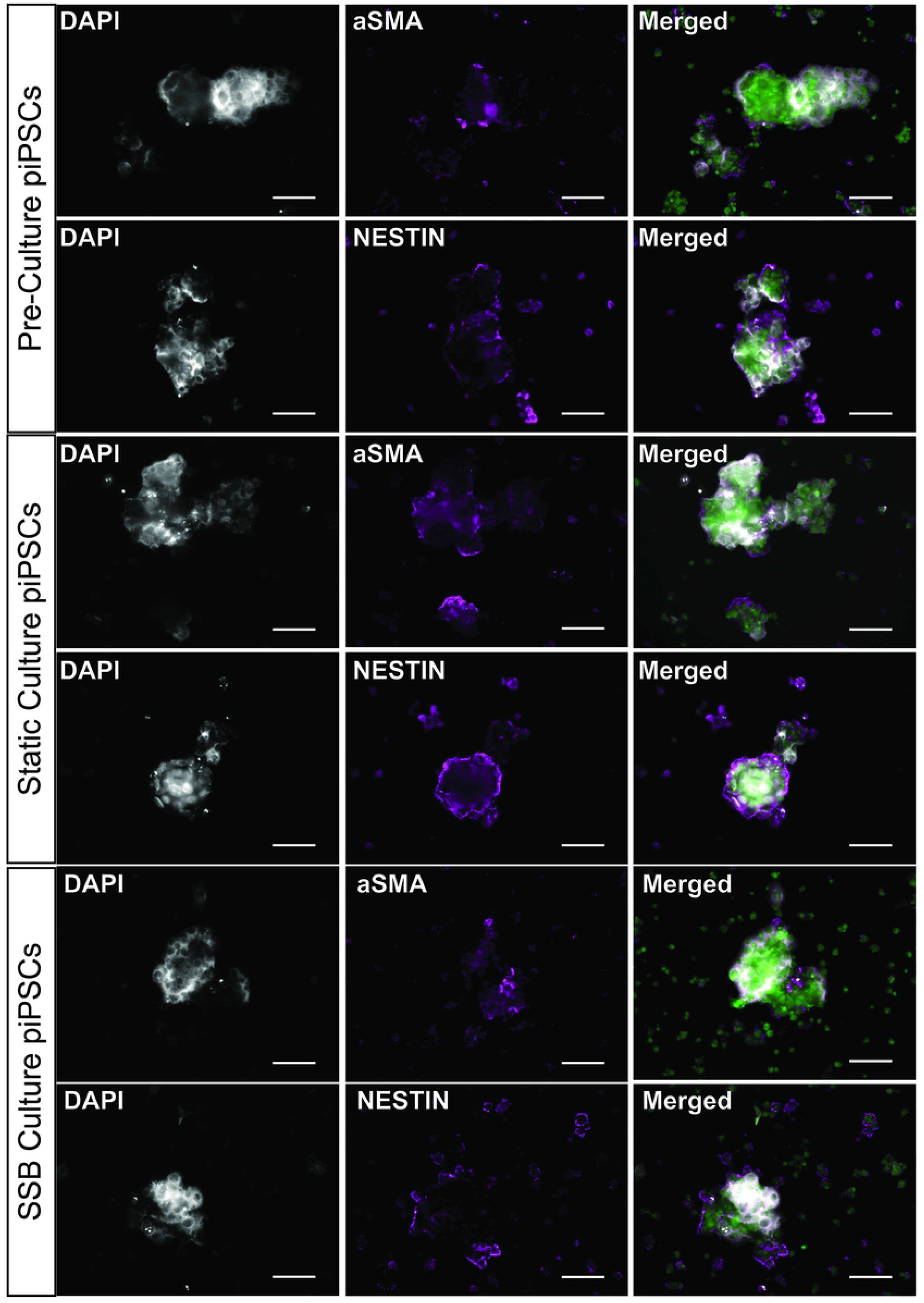
Differentiation-associated protein expression in day 3 spontaneously formed embryoid bodies from piPSCs. Intact embryoid bodies were collected and stained for ectoderm (NESTIN; purple) and mesoderm (*α*SMA; purple) proteins to illustrate differentiation into the respective germ lineages. Cells are constitutively expressing GFP (green). scale bar 50 μm.

### LIF-independent piPSCs cultured in SSB retain their pluripotency characteristics

Since the first porcine iPSC line examined in the current study relies on LIF to self-renew and proliferate, we sought to determine if a dermal fibroblasts-derived piPSC line that is LIF-independent would maintain iPSC key characteristics under the same dynamic environmental conditions. Applying SSB culture to this cell line addresses the question whether a possible interaction exists between the signaling pathways triggered by the shear stress and added growth factors that would affect piPSCs self-renewal and pluripotency. Similar to results obtained with the LIF-dependent piPSCs, SSB culture of the newly derived LIF-independent piPSCs (Fig. S2B) resulted in viability (Fig. 7A) and doubling times (Fig. 7B) comparable to those of the static culture (Fig. S2B) after 4 passages and cells maintained a stable karyotype (Fig.7C).

**Fig 7:**
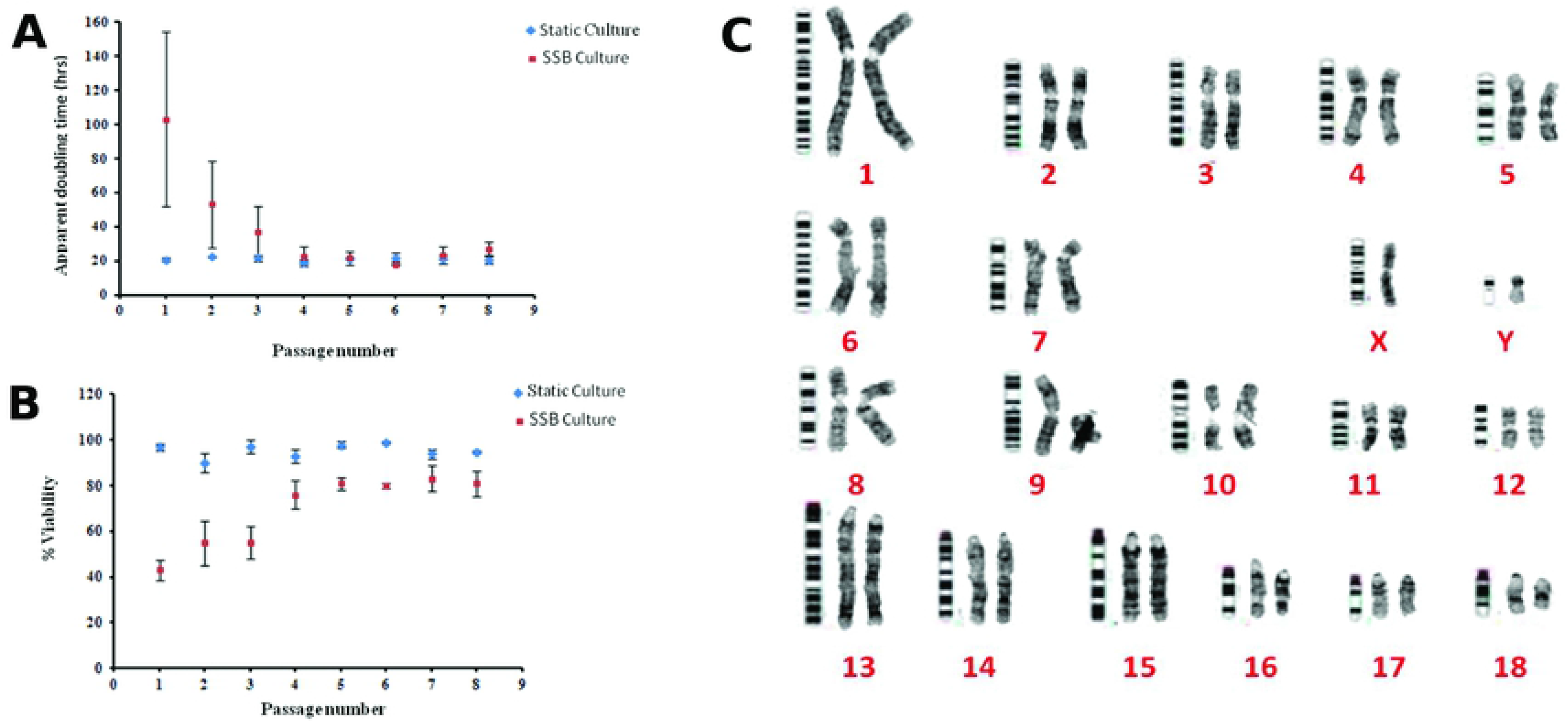
Apparent doubling time, viability and cytogenetic analysis. Apparent doubling (A) and cell viability (B) comparing SSB to static cultures over 8 passages. Cells were passaged every 3 days, and cells were collected at day 3 for analysis n=3. Error bars=SD. Cytogenetic analysis of piPSCs (C) image shows piPSCs from SSB culture. Cytogenetic analysis was performed on preculture piPSCs, P8 SSB piPSCs, and P8 static piPSCs.

Assessing the expression of pluripotency-associated genes, showed a down-regulation in *pSOX2* and *cMYC* gene expression in SSB culture as compared to static culture, while no significant difference in the expression of *OCT4, cMYC, KLF4* or *NANOG* was observed between static and SSB cultures (Fig. S2C). In contrast, immunocytochemistry did not detect a difference between SSB and static culture in the expression of exogenous hSOX2, OCT4, and SSEA-4, or Ki67 used here to assess cell proliferation (Fig. S2D). These observations further support the dependency of porcine iPSCs on the exogenous reprogramming genes to maintain their pluripotency.

To fully explore the potential of this newly generated LIF-independent piPSC line to differentiate into the three germ layers, we performed *in vitro* and *in vivo* differentiation assays. When compared to undifferentiated piPSCs, both static and SSB cultures-derived EBs (Fig. S3A) showed an up-regulation of *ACTA2, SOX17* and *NESTIN* expression, while they maintained *GATA4* (Fig. S3B). The gene expression of *NESTIN* differed in EBs between the two culture conditions. There was no noticeable difference in the expression of αSMA, GATA4 or NESTIN observed between the static and SSB-derived EBs (Fig. S3C). Surprisingly, EBs derived from static and SSB cultures maintained the expression of pluripotency markers *OCT4, cMYC* and *KLF4* during the course of this experiment (Fig. S3D), and further showed a significant increase in porcine *SOX2* expression, as compared to undifferentiated cells (Fig. S3D).

We then performed teratoma assays by injecting piPSCs cultured under static and SSB conditions subcutaneously to NOD-SCID mice. Histology analysis showed the presence of mesodermal, endodermal and ectodermal structures within the tissue formed from both static culture piPSCs (Fig. S4A) and SSB culture piPSCs (Fig. S4B). Expression of NESTIN and β III TUBULIN (as ectoderm markers), αSMA (as mesoderm marker) and GATA4 and α FETOPROTEIN (AFP) (as endoderm markers) was assessed and confirmed the differentiation of piPSCs into ectoderm (Fig. S4C), and mesoderm (Fig. S4C) but not endoderm based on the staining for GATA4 or AFP. To what extend the type of reprogramming strategy (lentiviral reprogramming *vs.* DOX-inducible transgene activation) affects the differentiation potential of piPSC deserves further investigation.

## Discussion

In the present work, we aimed to determine whether SSBs offered an efficient alternative culture platform to the two-dimensional static culture system for the expansion of piPSCs. Here we provide evidence that piPSC, similar to mESCs, can be expanded in SSBs using a shear force of 3.0 dyn/cm^2^ while maintaining their pluripotency potential.

Our study demonstrated that LIF-dependent piPSCs cultured in SSBs showed no difference in cell viability or apparent doubling time after the first passage, when compared to static culture. With the exception of P7 from static culture, which may have been due to an issue during piPSC dissociation and/or collection, the percent viability ranged between 90 and 95% for both static and SSB cultures. Our findings are consistent with previous studies using either similar or different SSB culture systems. Human iPSCs (hiPSCs) cultured in a pendulum-stirred system showed > 95% viability (21), whereas a study employing a triangular impeller and glass-etched baffles achieved a viability between 85 – 95% (10; 22). Two murine iPSC lines cultured in SSBs with an impeller similar to a magnetic stirrer bar that resembles the one used in the current study, also showed viability higher than 85% and 93% (13).

The proliferation of piPSCs in static and SSB cultures as assessed by Ki67 expression further validated the apparent doubling time data. For the LIF-dependent piPSCs, with the exception of the first passage, no difference was observed between the two types of cultures, which was consistent with previous work on human cells (23). Similarly, viability of the LIF-independent piPSCs increased with subsequent passages and was comparable to that observed in static culture. These data suggest that piPSCs can adapt rapidly to SSB culture, possibly via selection for a subpopulation able to survive under these conditions. Our results contrast, however, previous work on hiPSCs (12) and mESCs (8), where cell yield in SSB increased or decreased when compared to static controls, respectively. It is worth noting that optimization was required in both studies to circumvent the reduction of SSB-cultured cell yield compared to static culture, such as increasing agitation rate or adding supplements to culture medium. We speculate that the differences between our study and the aforementioned studies may be due to species specific differences, inoculation density, agitation rate or cell-specific responses to shear and the dynamic environment.

It is important to note that maintenance of the two piPSCs lines under dynamic conditions for an extended period of time had no impact on chromosomes stability. The disomy of the Y chromosome observed in the karyotype of LIF-dependent piPSC was already apparent in the starting cell population, which appeared to have arisen prior to the onset of the current experiments.

Expanding the LIF-dependent piPSCs line under dynamic conditions did not affect their pluripotency potential, as clearly confirmed by the sustained expression of OCT4, SOX2, cMYC and SSEA-4, which is consistent with previous reports on SSB culture of miPSCs (13) and hiPSCs (12). These piPSCs resemble hESCs in the expression of SSEA-4 and SSEA-1 (23), presumably due to the presence of LIF. piPSCs grown in the presence of LIF express SSEA-4 while piPSCs grown in the presence of FGF express SSEA-1 (24; 25; 26). This was further supported by the low expression of SSEA-4 observed in the LIF-independent piPSC line used in the current study. However, down-regulation of SOX2 and cMYC was observed in the LIF-independent piPSCs cultured in SSBs as compared to static conditions. SOX2 controlled the proliferation of adult stem cells (ASC) by regulating the expression of cMYC (27). Although speculative, it is possible that the piPSCs used in the current study regulate the expression of SOX2 to maintain their stemness. As Golden et al. (2017) reported, ESCs regulate the levels of SOX2 by activating a negative feedback loop via Akt signaling to retain their stem cell properties (28). Whether Akt signaling can be triggered as a result of the expression level of SOX2 alone or combined with other shear force-induced effects deserves further investigation, as a drastic change in morphology and function of human endothelial cells due to fluid shear stress-induced Akt phosphorylation has been reported (29).

One of the hallmarks of pluripotent cells is their ability to differentiate into three germ layers. Spontaneously formed EBs from pre-, static and SSB culture of LIF-dependent piPSCs expressed proteins representative of ectoderm, mesoderm and endoderm. NESTIN represents a reliable marker for neural progenitor cell differentiation from porcine cells (30; 31; 32). *ACTA2*, the gene encoding for the protein αSMA, was selected as a reliable marker for mesodermal differentiation for piPSCs (26), as well as hESC differentiation towards smooth muscle cells (33). *GATA6* has been shown in previous studies to be expressed upon endodermal lineage differentiation of piPSCs (30; 31) and is considered to play an integral role in the formation of human definitive endoderm from pluripotent stem cells (34). Our findings showed gene expression corresponding with ectoderm *(NESTIN)*, mesoderm *(ACTA2)*, and endoderm *(GATA6)* differentiation within 3 days of spontaneous embryoid body differentiation in cells cultured in both conventional adherent culture conditions and SSB. As expected, with the removal of DOX pluripotency-associated gene expression was down-regulated, offering further evidence of the differentiation of the piPSCs towards the germ lineages.

Overall, SSB culture conditions maintained the differentiation potential of both piPSC lines tested in the current study. Some differences, however, were noted between the two piPSC lines regarding the expression of pluripotency markers during the differentiation process. Perhaps an intrinsic feature of the LIF-independent piPSC, there was an up-regulation of SOX2 along with the differentiation markers in both static and SSB-derived EBs. A two-fold up-regulation in SOX2 expression induced differentiation of ESC into cells that express ectoderm, mesoderm and trophectoderm but not endoderm (35), however, the LIF-independent piPSCs reported here differentiated into derivatives of all three germ layers *in vitro.*

Formation of teratomas in NOD-SCID mice is considered a gold standard of pluripotency. The LIF-dependent iPSC failed to form teratomas. In previous work, LIF-dependent piPSCs were only able to form teratomas upon administration of DOX via drinking water (36). Dependence on DOX is consistent with the fact that the pluripotent state of piPSCs still requires the persistence of exogenous genes to maintain pluripotency, possibly due to the inadequate activation of endogenous genes to maintain pluripotency (37). However, the newly derived LIF-independent iPSCs differentiated into ectoderm and mesoderm *in vivo*, and this potential was maintained in SSB culture.

Here we demonstrate that piPSCs grown in static and SSB culture are similar in their functional characteristics. The downstream application of using SSBs is the potential for scale up. Considering the experimental parameters set by this study, such as seeding density and culture duration, the theoretical cell yield can be calculated to illustrate the number of piPSCs generated if all piPSCs were seeded in SSBs after each passage (*Supplementary information*). Using 10^9^ cells as a “therapeutic” dose, 1.406 x10^6^ piPSCs can be inoculated into a 50 mL SSB and scaled up over the following 3 passages yielding a total cell number of ~5.91 × 10^9^ piPSCs if scaled up to 2 – 10 L SSBs (Table S1). If this theoretical experiment was carried out using the static 6-well tissue culture plates, a “therapeutic” cell yield would be achieved by the 3^rd^ passage, however 143 6-well tissue culture plates would be required. As this study illustrates that the piPSCs cultured statically or in SSBs are functionally equivalent, scale up would be a more feasible option using SSBs rather than static tissue culture plates for future studies.

## Conclusions

We present here proof-of-principle for stirred suspension bioreactor culture of piPSCs, providing a scalable method of propagating piPSCs *in vitro.*

## Acknowledgements

We thank Drs. T. Ezashi, B. Telugu and M. Roberts for the GFP^+^ porcine iPSCs.

## Author Contributions

**Kyle Burrell**

Roles: Conceptualization, Data curation, Formal analysis, Investigation, Methodology, Validation, Visualization, Writing – original draft, Writing – review & editing

**Rkia Dardari**

Roles: Conceptualization, Data curation, Formal analysis, Investigation, Methodology, Validation, Visualization, Writing – review & editing

**Taylor Goldsmith**

Roles: Formal analysis, Investigation, Writing – review & editing

**Derek Toms**

Roles: Conceptualization, Methodology, Supervision, Writing – review & editing

**Daniel A.F. Villagomez**

Roles: Formal analysis, Investigation, Writing – review & editing

**W. Allan King**

Roles: Resources, Writing – review & editing

**Mark Ungrin**

Roles: Writing – review & editing

**Franklin West**

Roles: Resources, Methodology, writing-review & editing

**Ina Dobrinski**

Roles: Conceptualization, Funding acquisition, Methodology, Project administration, Resources, Supervision, Writing – review & editing

